# Impaired emotion recognition in *Cntnap2*-deficient mice is associated with hyper-synchronous prefrontal cortex neuronal activity

**DOI:** 10.1101/2023.10.19.563055

**Authors:** Alok Nath Mohapatra, Renad Jabarin, Natali Ray, Shai Netser, Shlomo Wagner

## Abstract

Individuals diagnosed with autism spectrum disorder (ASD) show difficulty in recognizing emotions in others, a process termed emotion recognition. While human fMRI studies linked multiple brain areas to emotion recognition, the specific mechanisms underlying impaired emotion recognition in ASD are not clear, partially due to the lack of appropriate tests in animal models. Here, we employed an emotional state preference (ESP) task to show that *Cntnap2*-knockout (KO) mice, an established ASD model, do not distinguish between conspecifics according to their emotional state. We assessed brain-wide local-field potential (LFP) signals during various social behavior tasks and found that *Cntnap2*-KO mice exhibited higher LFP theta and gamma rhythmicity than did C57BL/6J mice, even at rest. Specifically, *Cntnap2*-KO mice showed increased theta coherence, especially between the prelimbic cortex (PrL) and the hypothalamic paraventricular nucleus, during social behavior. Moreover, we observed significantly increased Granger causality of theta rhythmicity between these two brain areas, across all types of social behavior. Finally, optogenetic stimulation of PrL pyramidal neurons in C57BL/6J mice impaired their social discrimination abilities, including ESP behavior. Together, these results suggest that increased activity of PrL pyramidal neurons and their augmented synchronization with specific brain regions are involved in the impaired emotion recognition exhibited by *Cntnap2*-KO mice.

## Introduction

Social cognition involves the perception and interpretation of social cues transmitted between individuals, processes crucial for the appropriate adaptation of a subject to its social environment ^1,2^. The ability to recognize the emotional state of other individuals, termed emotion recognition, is crucial for a wide range of prosocial behaviors, such as emotion contagion, empathy and helping behavior ^3,4^ and is known to be impaired in Individuals diagnosed with autism spectrum disorder (ASD) ^5–7^. While fMRI studies have supplied considerable information regarding active brain areas during tasks of emotion recognition ^6–10^, including frontal lobe and amygdalar regions ^11,12^, specific deficits in brain activity and neuronal network dynamics that lead to impaired emotion recognition in ASD remain elusive. One valuable tool for deciphering such deficits is animal models of ASD, which allow for invasive monitoring and manipulation of brain neural activity during social behavior. Yet, reliable behavioral tasks to assess emotion recognition in animal models were lacking until only recently.

Of late, we and others have demonstrated that mice can discriminate between conspecifics according to the conspecific’s emotional state, thus providing a tool for assessing emotion recognition in murine models of ASD ^13–15^. This observation led to the development of a new behavioral task, termed by us emotional state preference (ESP). This task is based on the ability to discriminate between two stimulus animals simultaneously presented to a subject mouse, one of which was emotionally aroused by a given manipulation. While Ferretti *et al*. showed that C57BL/6J mice preferred to investigate stressed and relieved conspecifics more than neutral stimulus animals ^13^, we demonstrated that C57BL/6J mice preferred to investigate a stimulus animal socially isolated for seven days over a group-housed stimulus animal ^14^.

We previously demonstrated that mice expressing the A350V-encoding mutation in the *Iqsec2* gene, a mutation associated with ASD in humans ^16^, showed a specific deficit in the ESP task ^14^. Here, we sought to further validate this observation in another ASD murine model, so as to assess the generality of this deficit. One of the most established murine models of ASD is the *Cntanp2*-knockout (KO) mouse line, which does not express a functional copy of the *Contactin-associated protein 2* gene ^17^. Previous studies showed that such mice exhibit a reduced tendency for social interaction and that this can be reversed by inhibiting the excitability of medial prefrontal cortex (mPFC) pyramidal neurons or by activating inhibitory GABAergic interneurons in this region ^18^. Natably, recognition of stress and relieved states by C57BL/6J mice was found to be dependent upon somatostatin-expressing GABA-ergic inhibitory interneurons in the mPFC ^15^. However, emotion recognition by *Cntnap2*-KO mice has yet to be examined.

Here, we addressed this issue by employing the ESP paradigm to examine emotion recognition in *Cntnap2*-KO mice. We found that these mice do not prefer to investigate aroused over neutral conspecifics. Surprisingly, we found that even wild-type (WT) offspring of *Cntnap2*^−/+^ mice were impaired in this behavior, suggesting that the etiology of this impairment involves not only a subject’s genotype, but also the genotype of its parents and/or littermates. We further simultaneously recorded local field potential (LFP) signals from multiple brain areas linked to social behavior and found hyperactive theta and gamma rhythmicity in the brains of *Cntnap2*-KO mice, as compared to C57BL/6J mice. Specifically, synchronization between the mPFC and several hypothalamic and amygdalar areas was consistently modified in *Cntnap2*-KO mice across multiple social discrimination tasks. Finally, using optogenetic stimulation, we demonstrated that stimulating mPFC pyramidal neurons at either theta or gamma frequency impaired the ability of C57BL/6J mice to discriminate between various types of conspecifics and specifically, between emotionally-aroused and neutral stimuli, similarly to *Cntnap2*-KO mice. These results suggest that the modified mPFC activity exhibited by *Cntnap2*-KO mice does not merely cause social avoidance, as previously suggested, but rather interferes with social recognition and discrimination, thus creating a complex deficit which seems to be highly related to ASD.

## Results

### *Cntnap2*-KO and WT littermates exhibit impaired emotion recognition

Since patients diagnosed with ASD are known to exhibit impaired emotion recognition ^3–5^, we first examined whether *Cntnap2*-KO mice are impaired in terms of ESP using two variations of the test that assesses this trait. Specifically, in the stress-state preference (SSP) task, subjects simultaneously encountered a stressed stimulus animal and a neutral animal, whereas in the isolation-state preference (ISP) task, subjects encountered a socially isolated stimulus animal and a group-housed animal. Each task consisted of a 5 min encounter session, which followed a 5 min baseline period during which time no stimulus was introduced into the arena. As a control, we conducted a social preference (SP) task in which the subjects encountered a novel animal stimulus and an object (Fig. 1A). In this task, both WT and *Cntnap2*^−/−^ (*Cntnap2*-KO) male littermates showed similar SP behavior, namely, investigating the stimulus animal for significantly more time than the object (Fig. 1D-E). Surprisingly, when conducting the SSP (Fig. 1B) and ISP (Fig. 1C) tasks with these same two genotypes, we found that both WT and KO mice did not discriminate between the two stimulus animals in either task (Fig. 1F-G, H-I). Since previous studies showed that behavioral deficits in ASD genetic models are sometimes exhibited by WT animals raised with mutant littermates by mutant parents ^19,20^, we generated “pure WT” mice by breeding two WT parents ^21^. We found that pure WT mice perform normally in the SSP and ISP tasks (Fig. 1J-K). These results show that *Cntnap2*-KO mice are impaired in terms of their ESP behavior, although the etiology of this impairment involves not only the subject’s genotype but also the genotypes of its parents and/or littermates. As pure WT animals cannot be littermates of KO animals and since the genetic background of the KO mice is that of the C57BL/6J (C57) mouse strain, we continued this study by comparing brain activity of KO and C57 mice during social behavior.

**Figure 1.**
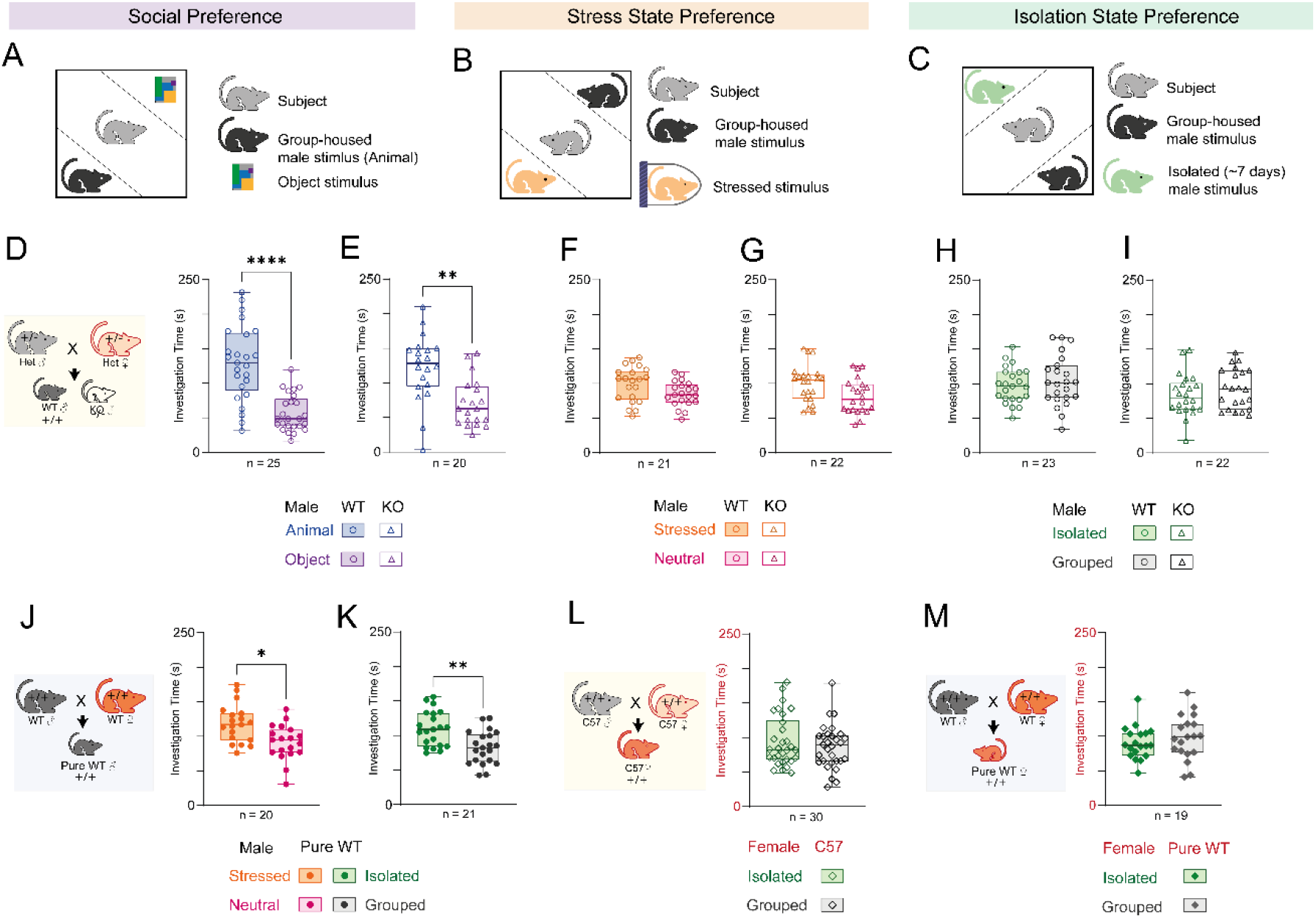
*Cntnap2*-KO and WT littermates exhibit impaired emotional state preference (ESP) behavior. **A.** Schematic representation of the social preference (SP) task. **B.** As in **A**, for the SSP task. **C.** As in **A**, for the ISP task. **D.** Median (in box plot*) time dedicated by WT littermates to investigating the animal (blue) or object (purple) stimulus during the 5 min encounter period of the SP task (paired t-test, n = 25 sessions, t (24) = 4.934, P<0.0001). The breeding scheme of WT and KO littermates is shown in the inset on the left. **E.** As in **D**, for KO littermates (Paired t-test, t (19) = 4.934, P = 0.0066). **F.** As in **D**, for SSP (t (20) = 1.869, P = 0.0763). **G.** As in **E**, for SSP (t (21) = 1.870, P = 0.0755). **H.** As in **D**, for ISP (t (22) = 0.5879, P = 0.5879). **I.** As in **E**, for ISP (t (21) = 0.8667, P = 0.3959). **J.** As in **F**, for pure WT animals produced by breeding two WT littermates (see the breeding scheme in the inset on the left) (t (18) = 2.403, P = 0.0273). **K.** As in **J**, for the ISP task (t (20) = 2.987, P = 0.0073). **L.** As in **K**, using C57BL/6J female mice (t (20) = 1.429, P = 0.1658). See the breeding scheme in the inset on the left. **M.** As in **K**, using pure WT female mice (t (19) = 0.836, P = 0.4141). See the breeding scheme in the inset on the left. Box plot represents 25 to 75 percentiles of the distribution, while the bold line is the median of the distribution. Whiskers represent the smallest and largest values in the distribution. *p<0.05, **p<0.01, ****p<0.0001, Paired t-test.

Accordingly, we employed four behavioral tasks, namely, the SP and the ISP tasks, a sex preference (SxP) task in which the subjects encountered a male and a female stimulus animal, and a free social interaction (FSI) task involving a same-sex, age-matched novel stimulus animal. For these experiments, we only used male subjects, as both C57 and pure WT female mice did not show a preference in the ISP task, which seems to be sex-specific (Fig. 1L-M). A multi-electrode array was implanted in the brain of each subject as previously described ^22^ and behavioral experiments were conducted 3 days later (see timeline in Fig. 2A). Each of the recorded subjects (11 C57 and 15 KO mice) performed up to three sessions of each task, as described in Fig. 2A, and the results from each session were separately analyzed.

**Figure 2.**
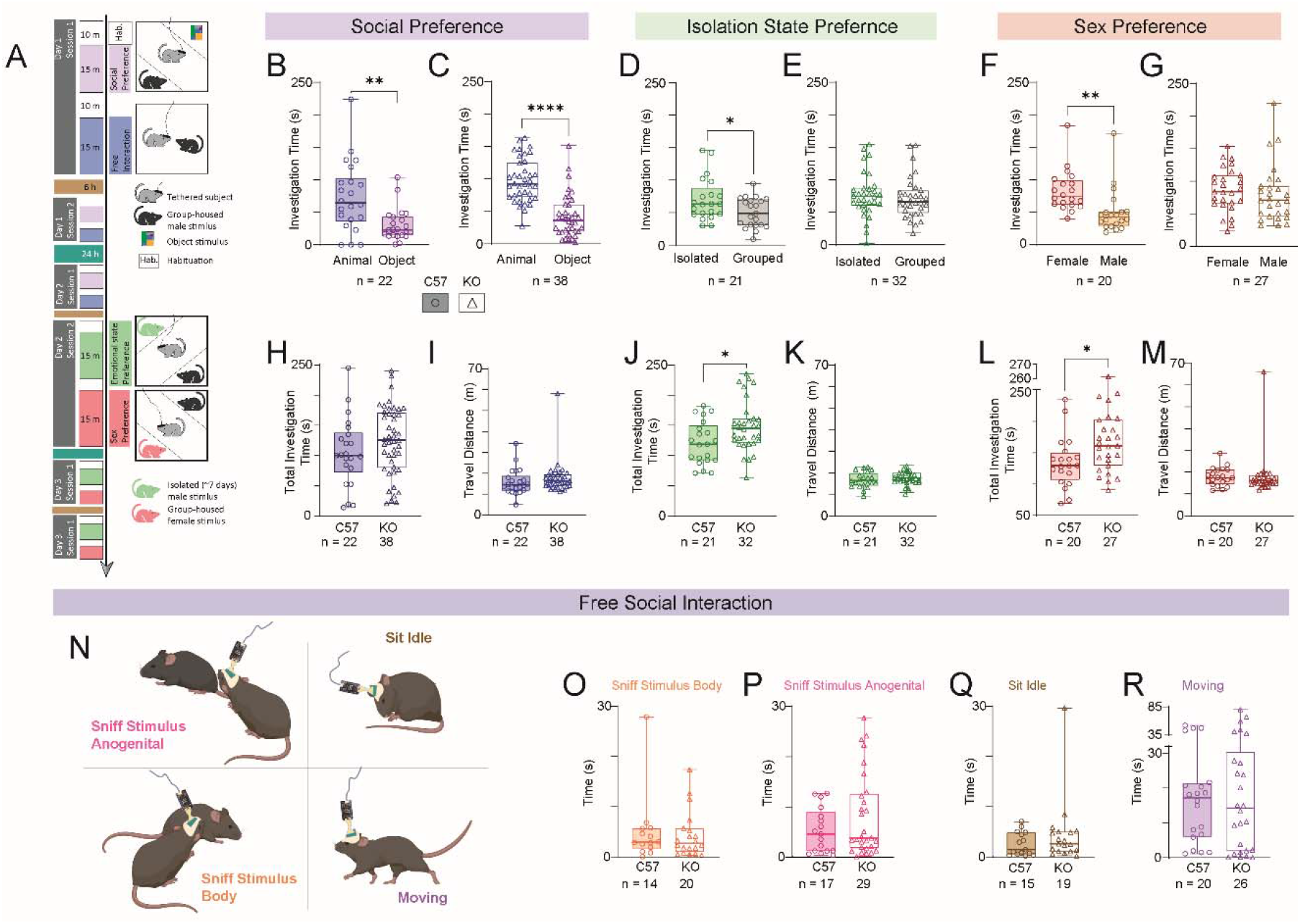
*Cntnap2*-KO male mice exhibit specific deficits in the ISP and SxP tasks. **A.** Timeline of the various tasks conducted by the electrode array-implanted mice. **B.** Median time dedicated by C57 mice to investigate the animal (blue) or object (purple) stimulus during the 5 min encounter period of the SP task (Wilcoxon matched-pairs signed rank test, W = −161, P = 0.0074). **C.** As in **B**, for KO mice (W = −639, P < 0.0001). **D.** As in **B**, for the ISP task (W = −121, P = 0.0351). **E.** As in **C**, for the ISP task (W = −152, P = 0.16). **F.** As in **B**, for the SxP task (W = −146, P = 0.0049). **G.** As in **C**, for the SxP task (W = −80, P = 0.3483). **H.** Median time dedicated by C57 (left bar) and KO (right bar) mice to investigate both stimuli (total investigation time) during the 5 min encounter period of the SP task (Unpaired t test, t (65) = 1.265, P = 0.2103). **I.** As in **H**, for the distance traveled by the subjects during the SP task (Mann-Whitney test, U = 319, P = 0.1316). **J.** As in **H**, for the ISP task (t (51) = 2.439, P = 0.0159). **K.** As in **I**, for the ISP task (U = 267, P = 0.2149). **L.** As in **H**, for the SxP task (t (45) = 2.458, P = 0.0179). **M.** As in **I**, for the SxP task (U = 224, P = 0.2484). **N.** Schematic representations of the four poses analyzed using DeepLabCut + SimBA analyses. **O.** Median time dedicated by C57 (left bar) and KO (right bar) mice to sniff the body of the stimulus animal during the 5 min encounter period of the FSI task (Mann Whitney test, U = 128, P = 0.6913). **P.** As in **O**, for sniffing the anogenital region of the stimulus animal (U = 211, P = 0.4297). **Q.** As in **O**, for sitting idle (U = 112, P = 0.3024). **R.** As in **O**, for moving (U = 250, P = 0.8349). Box plot represents 25 to 75 percentiles of the distribution, while the bold line is the median of the distribution. Whiskers represent the smallest and largest values in the distribution. *p<0.05, **p<0.01, ****p<0.0001, Unpaired t test, Mann-Whitney test or Wilcoxon matched-pairs signed rank test. See Fig. S1 for more details regarding the SimBA model.

We found that both genotypes (C57 and KO) showed a significant preference to investigate the social stimulus over the object in the SP task (Fig. 2B-C). However, while C57 mice also showed a preference to investigate the isolated stimulus animal in the ISP task (Fig. 2D) and the female stimulus animal in the SxP task (Fig. 2F), KO mice did not show any preferences in the two tasks (Fig. 2E, G). Interestingly, while there was no significant difference between the two genotypes in terms of the mean distance traveled in the arena during the encounter period of any task (Fig. 2I, K, M), KO mice displayed significantly longer total investigation times (of both stimuli together) in the ISP and SxP tasks (Fig. 2H, J, L). This suggests that KO mice are generally more exploratory than are C57 mice.

In the last task (i.e., the FSI task), each subject was exposed to a novel stimulus animal for five minutes in the empty arena, and the behavior of the subject was tracked using DeepLabCut (DLC) software ^23^, and analyzed by SimBA behavioral segmentation analysis (Fig. 2N and Fig. S1) ^24^. We found no differences between the two genotypes in any of the behavioral variables analyzed (Fig. 2O-R), other than the number of “sitting idle” events (Fig. S1E). Thus, we identified deficits in the social behavior of KO mice, specifically in the ISP and SxP tasks.

### *Cntnap2*-KO mice display augmented theta and gamma rhythmicity

To assess brain activity during behavior, we recorded local field potential (LFP) signals from microelectrode array-implanted mice while they performed various behavioral tasks (Fig. 3A). The location of each electrode tip was verified post-mortem in each mouse (Fig. S1K) and only regions where an adequate sample size was available for both C57 and KO mice were considered. Of all recorded brain areas (nine common to C57 and KO mice), we analyzed signals from seven social behavior-associated brain regions (see Methods, supp. Table 1). These brain areas addressed included the anterodorsal part of the medial amygdala (MeAD), the nucleus accumbens core (AsbC) and shell (AcbSh), the prelimbic (PrL) and infralimbic (IL) cortices, the hypothalamic paraventricular nucleus (PVN) and the lateral septum (LS) (Fig. 3B).

**Figure 3.**
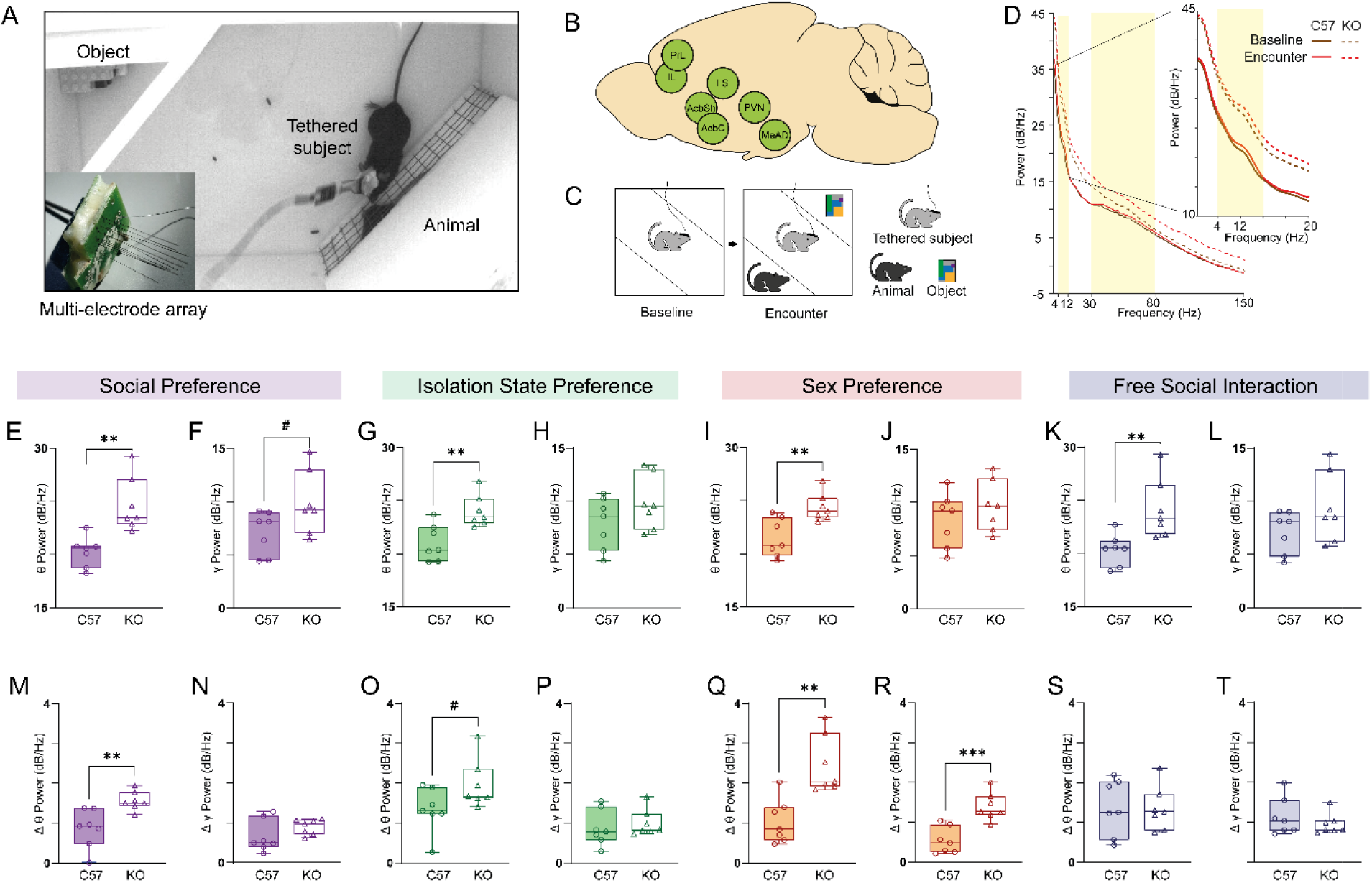
*Cntnap2*-KO mice exhibit higher levels of theta and gamma power in all tasks. **A.** A picture of a recorded subject mouse in the arena during the SP task. Inset - a picture of a microelectrode array. **B.** Schematic representation of the seven recorded brain areas analyzed. **C.** Schematic representation of the two stages of a recording session (SP, in this example), i.e., baseline and encounter. **D.** Example plotted power spectral density (PSD) profiles for LFP signals recorded from C57 (continuous lines) and KO (dashed lines) subject mice during the baseline (purple) and encounter (red) periods of a single SP session. Yellow areas denote the theta (4-12 Hz) and gamma (30-80 Hz) bands. Inset – the same curves showing the theta range at higher resolution. **E.** Median theta power of LFP signals recorded during the baseline period of all SP experiments, across all seven brain regions, for C57 (left bar) and KO (right bar) subjects (Unpaired t test, n = 7 Brain regions, t (12) = 4.036, P = 0.0016). **F.** As in **E**, for gamma power (t (12) = 2.053, P = 0.0626). **G-H.** As in **E-F**, for ISP sessions (**G**: t (12) = 3.476, P = 0.0046; **H**: t (12) = 1.437, P = 0.176). **I-J.** As in **E-F**, for SxP session (**I**: t (12) = 3.457, P = 0.0047; **J**: t (12) = 0.9996, P = 0.3372). **K-L.** As in **E-F**, for FSI session (**K**: t (12) = 3.204, P = 0.0076; **L**: t (12) = 1.587, P = 0.1385). **M-T.** As in **E-L**, for Δpower during the encounter, as compared to baseline. **M**: t (12) = 3.48, P = 0.0045; **N**: t (12) = 1.437, P = 0.1726. **O**: t (12) = 2.024, P = 0.0658; **P**: t (12) = 0.59, P = 0.56. **Q**: t (12) = 4.101, P = 0.0015; **R**: t (12) = 4.704, P = 0.0005. **S**: t (12) = 0.081, P = 0.9363; **T**: t (12) = 0.9117, P = 0.3799. Box plot represents 25 to 75 percentiles of the distribution, while the bold line is the median of the distribution. Whiskers represent the smallest and largest values in the distribution. #p<0.066, *p<0.05, **p<0.01, ****p<0.0001, Unpaired t test.

We first analyzed the mean power of theta and gamma rhythmicity of LFP signals recorded during the 5 min-long baseline period, during which time there were no stimuli in the arena, across all the brain regions listed above (Fig. 3C-D). We measured higher theta power in KO mice, as compared to C57 mice, in all four tasks, with this difference being statistically significant in all cases (Fig. 3E, G, I, K). Gamma power showed a similar trend, albeit without statistical significance (Fig 3F, H, J, L). These differences between the genotypes were observed even when we considered only the first session for each mouse, thus excluding the possibility that the changes were associated with expectation of a social encounter (Fig. S1L-M). The results suggest that KO mice exhibit a higher level of a certain internal state even before the beginning of the behavioral task.

We next analyzed the mean change (Δ) in theta and gamma powers recorded during the 5 min-long encounter, as compared to baseline power values (Fig. 3D). In all cases, LFP power during the encounter was higher than at baseline, as reflected by positive Δpower values in both the theta and gamma bands (Fig. 3O-V). However, the KO mice exhibited significantly higher increases in theta rhythmicity in all three social discrimination tasks (Fig. 3M, O, Q), while the gamma power change was only significantly higher in the SxP task (Fig. 3N, P, R). In contrast to the discrimination tasks, we did not find a significant difference in power change between genotypes during the FSI task (Fig. 3S-T). Overall, these results suggest a higher internal state, reflected by stronger theta rhythmicity, in KO mice than in C57 mice, which was further enhanced during the encounter stage of the various social discrimination tasks.

### *Cntnap2*-KO mice exhibit hyper-synchronized brain activity during social interaction

While the power of LFP rhythmicity may reflect the internal state of the animals ^25,26^, coherence between brain regions is thought to reflect functional connectivity between them ^27^. When analyzing the coherence of LFP rhythmicity across all pairs of brain regions considered, we found that the mean coherence did not differ between tasks and genotypes during the baseline period for both theta and gamma rhythms (Fig. 4A, C). These results suggest that the mean coherence did not reflect the initial internal state that affected the mean LFP power (Fig. 3E-L). However, the normalized change in theta coherence during the encounter showed a significant difference (after correcting for multiple comparisons) between genotypes, with KO animals showing higher coherence changes in all tasks, other than the SP task (Fig. 4B). No significant differences were observed for the change in gamma coherence (Fig. 4D). Thus, theta rhythmicity during the various tasks seems to be hyper-synchronized across the recorded brain regions in KO mice. Notably, the coherence between the PrL and PVN seemed to be especially high in KO animals, as compared to C57 mice, for both theta and gamma rhythmicity (Fig. 4B, D, filled circles; see also heat-maps in Fig. 4E-L). We, therefore, analyzed the coherence between this pair of brain regions in each task separately and found a significantly higher encounter-induced change in theta coherence in KO mice in all cases (Fig. 4M-P). At the same time, changes in gamma coherence only showed a significant difference in the SP and SxP tasks (Fig. 4Q, S). These results suggest higher synchronization in theta rhythmicity of KO mice during social behavior, especially between the PrL and PVN.

**Figure 4.**
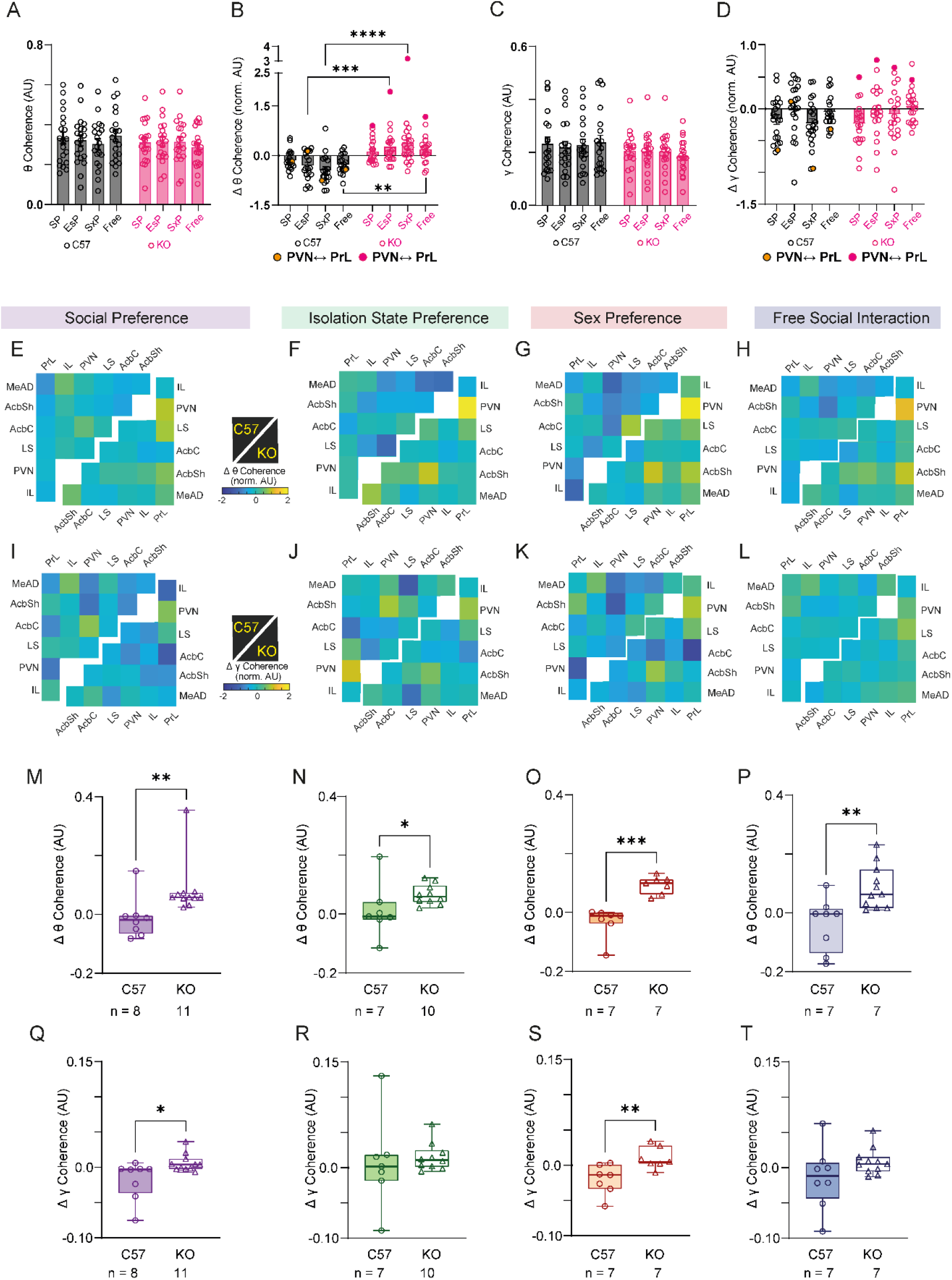
*Cntnap2*-KO mice show higher theta coherence during the various tasks, especially between the PrL and PVN. **A.** Mean (±SEM) theta coherence across all brain region pairs (n= 20 for each task) during the baseline of all tasks in C57 (black bars) and KO (red bars) subjects (Two-way ANOVA. Genotype: F (1, 152) = 1.459, P = 0.3104; Tasks: F (3, 152) = 0.1049, P = 0.9571; Interaction: F (3, 152) = 0.6683, P = 0.5727). **B.** Mean (±SEM) normalized change in theta coherence across all pairs of brain regions (n= 20 for each task) during the encounter period, as compared to baseline, of all tasks conducted by C57 (black bars) and KO (red bars) subjects. The data points representing the coherence between the PrL and PVN are denoted as filled dots. (2-way ANOVA. Genotype: F (1, 152) = 52.27, P <0.0001; Tasks: F (3, 152) = 0.04024, P = 0.9892; Interaction: F (3, 152) = 2.496, P = 0.062) **C-D.** As in **A-B**, for gamma coherence. **C:** Genotype: F (1, 152) = 4.081, P = 0.0451; Tasks: F (3, 152) = 0.061, P = 0.98; Interaction: F (3, 152) = 0.3091, P = 0.8188). **D:** Genotype: F (1, 152) = 0.4696, P = 0.4942; Tasks: F (3, 152) = 3.260, P = 0.0232; Interaction: F (3, 152) = 0.9164, P = 0.4346. **E.** Heat-map matrices of the normalized change in theta coherence during the SP task across all recorded pairs of brain regions for C57 (upper right matrix) and KO mice (lower left matrix). Note that white spots represent a brain region pairs with an inadequate sample size. **F.** As in **E**, for ISP sessions. **G.** As in **E**, for SxP sessions. **H.** As in **E**, for FSI sessions. **I-L.** As in **E-H**, for the normalized change in gamma coherence. **M.** Median change in theta coherence between the PrL and PVN across all SP sessions for the C57 (circles, left) and KO (triangles, right) subjects (Mann Whitney test, U = 10, P = 0.0036). **N.** As in **M**, for ISP sessions (U = 12, P = 0.025). **O.** As in **M**, for SxP sessions (U = 0, P = 0.0006). **P.** As in **M**, for FSI session (U = 10, P = 0.0036). **Q-T.** As in **M-P**, for the change in gamma coherence. **Q:** U = 14, P = 0.0121; **R:** ISP: U = 26, P = 0.4173; **S:** SxP: U = 4, P = 0.007; **T:** FSI: U = 4, P = 0.1518. Box plot represents 25 to 75 percentiles of the distribution, while the bold line is the median of the distribution. Whiskers represent the smallest and largest values in the distribution. **A-D**: **p<0.01, ***p<0.001, ****p<0.0001 Šídák’s multiple comparison test after two-way ANOVA; **M-T:** *p<0.05, **p<0.01, ***p<0.001, Mann Whitney test.

### The prelimbic cortex in *Cntnap2*-KO mice shows modified synchronization during social interaction

To further explore the possibility that KO mice exhibit modified synchronization between brain region, we analyzed Granger’s causality (GC), a measure which assesses predictability of the rhythmicity in one region according to the rhythmicity in another region, for a given pair of brain regions. The GC was separately calculated for each direction of the possible interaction between each pair of brain regions considered in this study. Interestingly, the GC level in the PrL to PVN direction in the theta band was consistently higher in KO mice than in C57 animals, with the difference being statistically significant (after correcting for multiple comparisons) across all tasks, other than the SxP task (Fig. 5 A-D). In the gamma band, we found a consistent and significantly lower GC in the PrL to MeAD direction in KO mice across all tasks (Fig. 5 E-H). Following this screen, we compared GC values between PrL and PVN in each direction across all sessions. We found a significant difference between C57 and KO animals in the theta GC in the PrL to PVN direction in all tasks, while in the other direction (i.e., PVN to PrL), a significant difference was only seen in the FSI task (Fig. 5I-P). Similarly, a significant difference was found in the PrL to MeAD direction for gamma GC values across all tasks, while in the other direction, such a difference was only noted in the SP task (Fig. 5Q-X). Overall, these results suggest that the PrL of KO animals exhibit hyper-synchronous neural activity with several other brain regions during social behavior.

**Figure 5.**
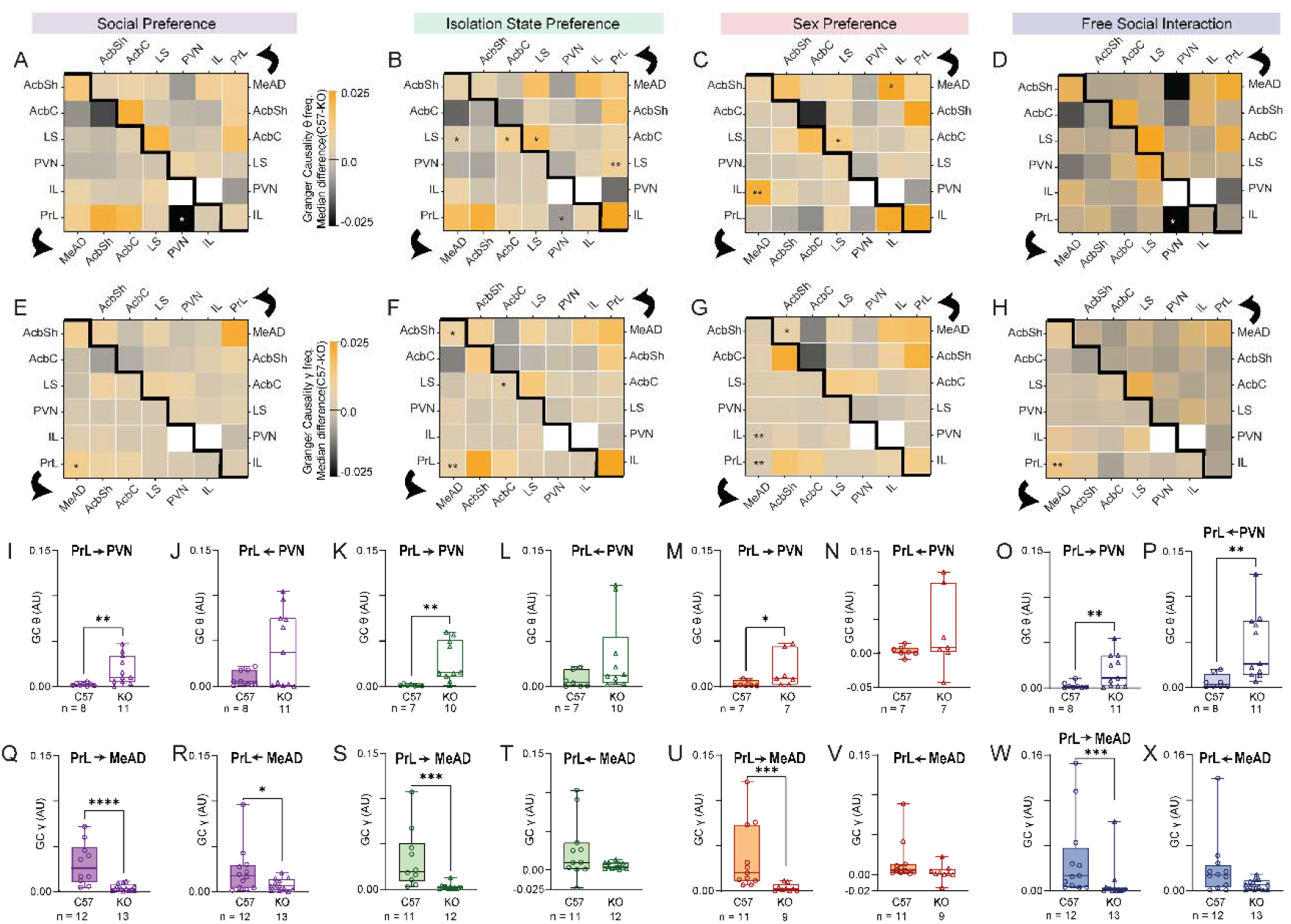
Theta GC values from the PrL to specific areas consistently differ between KO and C57 subjects. **A.** Matrix heat-map of the theta band GC differences between C57 and KO mice across all brain region pairs in a SP task session. Note that the lower left matrix represents the GC in one direction, as denoted by the black arrow, while the upper right matrix represents the other direction. White spots represent pairs with inadequate sample size. **B.** As in **A**, for ISP sessions. **C.** As in **A**, for SxP sessions. **D.** As in **A**, for FSI sessions. **E-H.** As in **A-D**, for gamma band GC differences. **I.** Median theta band GC values from the PrL to the PVN across all SP sessions for C57 (left bar) and KO (right bar) subjects (Mann Whitney test, U = 8, P = 0.0018). **J.** As in **I**, for theta band GC values from the PVN to the PrL (U = 34, P = 0.4421). **K-L.** As in **I-J**, for ISP sessions (**K:** U = 5, P = 0.002; **L:** U = 18, P = 0.1088) **M-N.** as in **I-J**, for SxP sessions (**M:** U = 5, P = 0.0111; **N:** U = 23, P = 0.9). **O-P.** As in **I-J**, for FSI sessions (**O:** U = 12, P = 0.0068; **P:** U = 8, P = 0.0018). **Q-X.** As in **I-P**, for gamma GC. **Q:** U = 7, P < 0.0001; **R:** U = 46, P = 0.0398; **S:** U = 33, P = 0.0439; **T:** U = 51, P = 0.38; **U:** U = 4, P = 0.0001; **V:** U = 25, P = 0.067; **W:** U = 15, P = 0.0003; **X:** U = 4, P = 0.0678. Box plot represents 25 to 75 percentiles of the distribution, while the bold line is the median of the distribution. Whiskers represent the smallest and largest values in the distribution. **A-H**: *p<0.05, **p<0.01 FDR corrected Mann Whitney test; **I-X**: *p<0.05, **p<0.01, ***p<0.001, Mann Whitney test.

### Optogenetic stimulation of PrL pyramidal neurons abolishes emotional state preference

The data presented thus far point to the PrL in KO mice as consistently showing higher changes in theta power and synchronization with other regions during social behavior. We, therefore, hypothesized that the hyper-active theta rhythmicity in the PrL could reflect hyper-excitability of PrL pyramidal neurons, which may elicit the behavioral impairments observed in the KO mice. To further examine this possibility, we examined the effect of stimulating PrL pyramidal neurons during the various discrimination tasks. Accordingly, we used the AAV viral vector to transfect CamK2a-postive neurons in the PrL cortex (presumably pyramidal neurons) of both C57 and KO animals (n = 6 animals per group) with Channelrhodosin-2 (ChR2) (Fig. 6A-B). We then used an optic fiber implanted into the PrL (Fig. S2A) to apply optogenetic stimulation at either 10 or 30 Hz (or applied no stimulation at all) (Fig. S2B-C) during each of the three social discrimination tasks, which were randomized over three days of experimentation (see timeline in Fig. 6B). Interestingly, we found that in all tasks, stimulation at 10 and at 30 Hz disrupted the ability of C57 mice to prefer one stimulus animal over the other (Fig. 6C-K). Moreover, both optogenetic stimulation protocols also disrupted the ability of KO mice to discriminate between the stimulus animal and object during the SP task (Fig. 6D-E), the only social discrimination task in which non-stimulated KO mice performed normally (Fig. 6C). In contrast, optogenetic stimulation caused no behavioral changes in KO mice during the SxP and ISP tasks (Fig. 6G-K). Furthermore, both optogenetic stimulation protocols resulted in identical behavior of C57 and KO mice in all three tasks (other than a slight change in the SxP test, Fig. 6D-K). These data suggest that hyper-activity of PrL pyramidal neurons at either theta or gamma frequency causes a lack of discrimination between stimulus animals in all three tasks in a manner similar, although not identical, to the impaired discrimination exhibited by KO animals.

**Figure 6.**
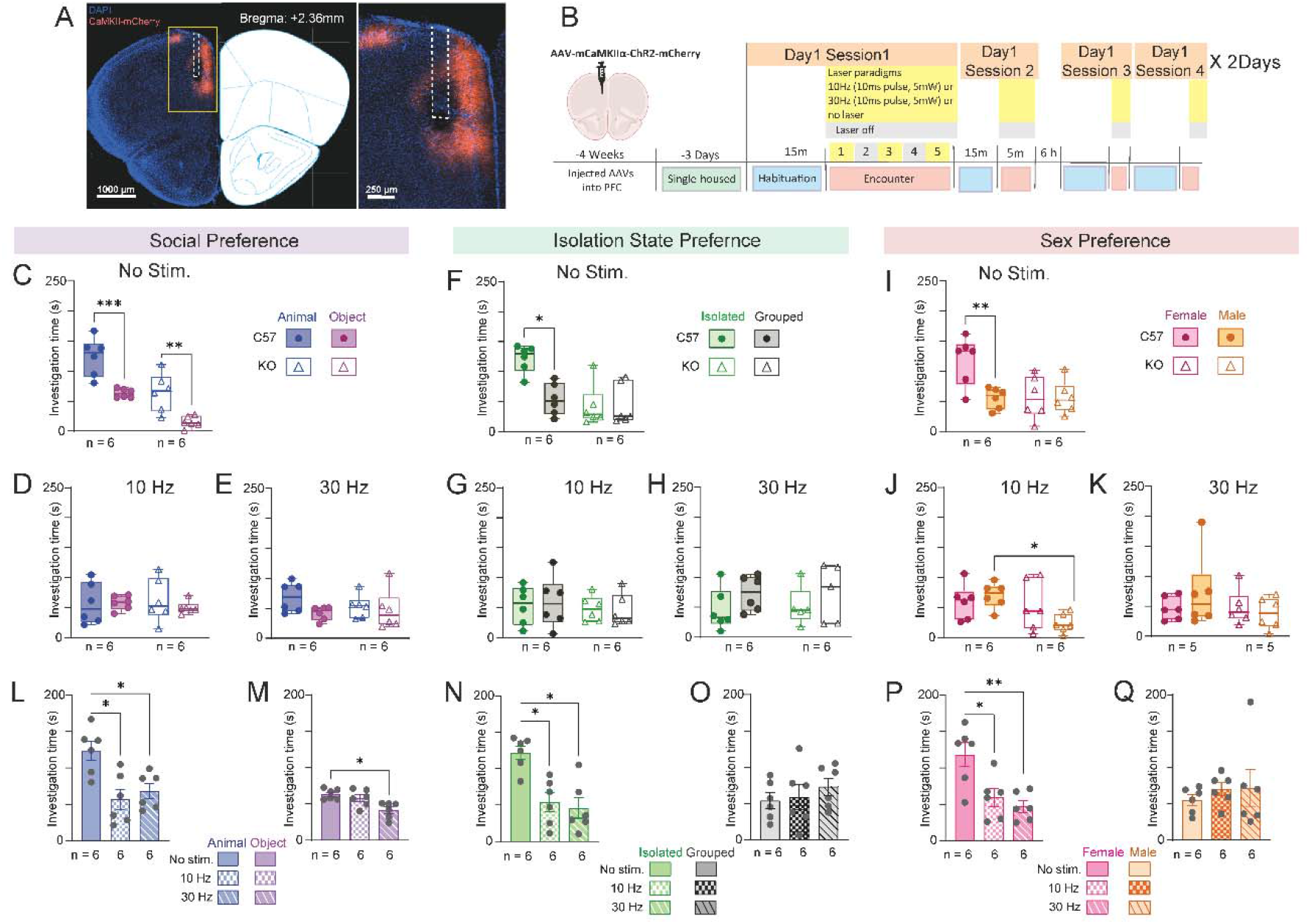
Optogenetic stimulation of PrL pyramidal neurons abolishes any preference for any given stimulus in all social discrimination tasks. **A.** A picture showing a coronal brain slice of the prefrontal cortex from a mouse injected with AAV virus carrying Chr2-mCherry under control of the CamK2a promoter on the left, and the corresponding mouse brain atlas slice on the right. The PrL area at higher spatial resolution is shown to the right of the brain atlas profile. **B.** The timeline of the various sessions conducted with the virus-injected mice. **C.** Median time dedicated by the virus-injected mice implanted with an optic fiber to investigate the animal stimulus (blue) or object (purple) during SP task sessions conducted by C57 (filled circles) and KO (empty triangles) mice without optogenetic stimulation (two-way RM ANOVA. Genotype: F (1, 10) = 29.3, P = 0.0003; Stimulus: F (1, 10) = 36.23, P = 0.0001; Interaction: F (1, 10) = 0.3894, P = 0.5466). **D.** As in **C**, with a 10 Hz optogenetic stimulation applied throughout the encounter stage (Genotype: F (1, 10) = 0.007, P = 0.9; Stimulus: F (1, 10) = 0.2, P = 0.61; Interaction: F (1, 10) = 0.43, P = 0.52). **E.** As in **F**, for a 30 Hz stimulation (Genotype: F (1, 10) = 0.32, P = 0.57; Stimulus: F (1, 10) = 2.34, P = 0.15; Interaction: F (1, 10) = 0.936, P = 0.35). **F-H.** As in **C-E**, for ISP sessions. **F:** Genotype: F (1, 10) = 22.6, P = 0.0008; Stimulus: F (1, 10) = 5.123, P = 0.0471; Interaction: F (1, 10) = 5.818, P = 0.0366; **G:** Genotype: F (1, 10) = 0.67, P = 0.43; Stimulus: F (1, 10) = 0.004, P = 0.95; Interaction: F (1, 10) = 0.11, P = 0.75; **H:** Genotype: F (1, 9) = 0.039, P = 0.84; Stimulus: F (1, 9) = 2.381, P = 0.157; Interaction: F (1, 9) = 0.033, P = 0.859. **I-K.** As in **C-E**, for SxP sessions. **I:** Genotype: F (1, 10) = 4.226, P = 0.0669; Stimulus: F (1, 10) = 8.713, P = 0.0145; Interaction: F (1, 10) = 8.585, P = 0.015; **J:** Genotype: F (1, 10) = 7.16, P = 0.023; Stimulus: F (1, 10) = 0.487, P = 0.5; Interaction: F (1, 10) = 2.295, P = 0.16; **K:** Genotype: F (1,10) = 1.097, P = 0.31; Stimulus: F (1, 10) = 0.2516, P = 0.6268; Interaction: F (1, 10) = 1.197, P = 0.2996. **L.** Mean (±SEM) time dedicated by the virus-injected mice implanted with an optic fiber to investigate the animal stimulus during SP sessions by C57 subject mice across the three optogenetic stimulation protocols (RM ANOVA, F (1.668, 8.342) = 11.75, P = 0.0047). **M.** As in **L**, for the object stimulus (F (1.584, 7.92) = 7.504, P = 0.018). **N.** As in **L**, for the isolated stimulus animal during ISP sessions (F (1.58, 7.898) = 15.63, P = 0.018). **O.** As in **L**, for the group-housed stimulus animal during ISP sessions (F (1.734, 8.668) = 0.907, P = 0.4245). **P.** As in **L**, for the female stimulus animal during a SxP task session (F (2, 10) = 9.243, P = 0.0053). Box plot represents 25 to 75 percentiles of the distribution, while the bold line is the median of the distribution. Whiskers represent the smallest and largest values in the distribution. **C-K**: *p<0.05, **p<0.01, ***p<0.001 Šídák’s multiple comparison test after 2-way RM ANOVA; **L-Q**: Šídák’s multiple comparison test after RM ANOVA.

Finally, since previous studies suggested that stimulating mPFC pyramidal neurons causes general avoidance of social stimuli and overall reduction in the tendency of a subject to engage in social interaction ^28,29^, we examined which behavioral variable was changed upon optogenetic stimulation of PrL pyramidal neurons in C57 mice. We found that in all cases, investigation of the preferred stimulus animal was reduced to the level of the less-preferred one, with the latter level being mostly unaffected (Fig. 6L-Q). Since the SxP and ISP tasks involve discrimination between two stimulus animals, these results suggest that optogenetic stimulation did not cause general social avoidance, but rather the loss of behavioral preference (Fig. 6L, N, P). Such a specific effect may be caused by a reduction in the valence of the preferred stimulus, which reduces the subject’s motivation to explore the stimulus. However, since the animal stimulus preferred in the SP tasks was the same type as the less-preferred stimulus animals in the other two tasks (i.e., a novel male group-housed mouse), our results contradict the possibility that a change in the absolute valence of the stimulus animal had occurred. Instead, the findings suggest that optogenetic stimulation reduced the relative valence of the preferred stimulus, and hence, the subject’s motivation to explore that stimulus. We, therefore, concluded that PrL pyramidal neurons may be involved in the display of preference during a competition between two rewarding stimuli, rather than in controlling the absolute motivation to display a specific behavior.

## Discussion

In the present study, we showed that *Cntnap2*-KO animals are impaired in their ESP behavior. Moreover, we found that even WT offspring of *Cntnap2*^+/−^ animals are impaired in this behavioral task, unlike other social discrimination tasks (such as the SP task) which they perform normally. This result, together with similar results obtained with *Shank3*-deficient rats (Jabarin *et al*., in preparation) and our previously published study of *Iqsec2* A350V mice ^14^, suggest that ESP behavior is especially sensitive to ASD-related mutations in mice. This may reflect the subtle behavioral, hormonal and physiological changes that distinguish an emotionally aroused animal from a relaxed animal. The same logic may explain why individuals diagnosed with ASD display specific impairments in emotion recognition, which require them to perceive and interpret subtle motion-induced changes in the facial expressions or body language of others. Thus, the ESP paradigm may be a useful tool for deciphering brain mechanisms underlying ASD-associated atypical social behavior.

Our observation that WT offspring of *Cntnap2*+/− animals were impaired in the ESP behavioral task is in accordance with results of other tasks found with *Neuroligin3*-KO mice, another well-established murine model of ASD ^19,20^. Interestingly, both genes (*Cntanap2* and *Neuroligin3*) encode synaptic proteins and both mutations seems to be associated with a modified excitatory/inhibitory (E/I) balance in the brain ^18,20^. This interesting phenotype may be related to gene-environment interactions, especially as related to the effect of the gut microbiome and its spread between parents and among littermates ^21,30^.

We used multisite electrophysiological recording from behaving mice ^22^ to explore population neuronal activity in multiple social behavior-associated brain regions during social behavior at the systems level. One intriguing observation was that KO mice exhibited generally stronger theta and gamma rhythms at baseline, even before the beginning of social interactions. This is in accordance with previously published single-cells recordings from the mPFC showing high levels of neuronal activity by *Cntnap2*-KO mice even before stimulus presentation ^31^. Given that augmented theta and gamma rhythms are associated with internal states such as arousal and attention ^32–36^, our results suggest the existence of a high level of a certain internal state in KO mice, which is in accordance with multiple studies demonstrating hyper-activity in these mice ^17,18,21^. While we did not detect longer distance traveled by KO mice during the social discrimination tasks, we did observe higher levels of general investigation time, which may reflect a hyper-active or hyper-exploratory state.

Other than the fundamentally higher levels of theta and gamma rhythms, we found a higher level of social encounter-induced theta rhythmicity in KO mice, as compared to C57 mice. Moreover, we found a generally high level of theta coherence induced by social encounter in all tasks, suggesting that the various recorded brain regions are over-synchronized in KO mice during social behavior. Such globally high coherence among brain regions may cause behavioral abnormalities by masking social context-induced specific patterns of coherence between brain regions, which may be required for proper recognition of and appropriate responses to specific stimuli ^37^. This result is in agreement with a recent study that used fMRI and cFos expression analyses to demonstrate macroscale functional hyper-connectivity in *Cntnap2*-KO mice ^38^. It should, however, be noted that these authors reported reduced microscale functional connectivity among several of the brain regions recorded in the current study. This contradiction between the studies may arise from the distinct aspect of functional connectivity measured in each of them (i.e., LFP theta rhythmicity vs. BOLD fMRI signal). In any case, both studies are in accordance with other studies showing that modified synaptic connectivity in the mPFC of *Cntnap2*-KO mice induces modified rhythmic population activity and synchrony in this area ^39^, as well as altered prefrontal functional connectivity associated with common genetic variants of the humans *CANTNAP2* gene ^40^. Together, these studies, which are in agreement with human studies showing a mix of hyper- and hypo-connectivity in ASD individuals ^41^, support the hypothesis that the behavioral symptoms of ASD are caused by aberrant functional connectivity that may occur as a result of developmental events ^42^.

We found that the PrL and PVN showed consistent and significantly augmented social behavior-induced hyper-synchrony across all paradigms. These results are very interesting, considering that the PVN is the main source of oxytocin to forebrain areas, in general, and to the mPFC, in particular ^43^. Oxytocin is a neuropeptide produced solely in the hypothalamic supraoptic and paraventricular nuclei ^44^ and is well known for its role in regulating social behavior ^45,46^. Recent studies showed that oxytocin administration can alleviate social behavior deficits exhibited by *Cntnap2*-KO mice ^38,47^ and that oxytocin is crucial for murine ESP behavior ^13^. Thus, our study complements these earlier efforts by demonstrating modified synchronization between the mPFC and PVN in *Cntnap2*-KO mice during social behavior, thus establishing a functional link between these two regions in the context of emotion recognition in ASD.

To examine whether the behavioral deficits we observed in KO mice may be caused by hyper-excitability and hyper-synchrony of mPFC pyramidal neurons, we applied rhythmic optogenetic stimulation to excite these cells in C57BL/6J mice during social encounters. In agreement with previous studies employing a similar stimulation protocol with all PrL pyramidal neurons ^29^ or to specific neuronal populations innervating either the NAc ^48^ or the BLA ^28^, we found that such stimulation impaired the SP behavior of the stimulated animals. However, unlike these previous studies, we also demonstrated that the same manipulation also abolished any preference in the SxP and ISP tasks. Notably, in all three tasks (SP, SxP and ISP), preference abolishment was induced by reducing the time dedicated by the subject to investigate the preferred stimulus animal, whereas no change was observed in the time dedicated to investigating the less-preferred stimulus, be it an object or stimulus animal. This, despite the fact that the same type of stimulus animal served as the preferred stimulus in the SP task and as the less-preferred stimulus in the other two tasks. As such, stimulation-induced reduction in investigation time was neither a general social avoidance, as suggested by previous studies ^29,49^, nor stimulus-specific avoidance. Instead, it seems to be a valence-specific response ^28^, expressed as a reduction in the desire to interact with a preferred stimulus.

Our observation that rhythmic optogenetic stimulation of PrL pyramidal neurons abolished the subjects’ preference in all tasks, supports the idea that hyper-active and hyper-synchronous PrL neural activity, as observed by us in *Cntnap2*-KO mice, may underlie the impaired SxP and ISP behaviors exhibited by these animals. Our observation that these animals function normally in the SP task is most probably explained by the fact that the difference between the two stimuli (object vs. conspecific) is highest in this task, as compared to all other tasks (two conspecifics). This evident difference makes SP the least challenging of all tasks, hence causes its resilience to the naturally-induced hyper-synchronous activity of PrL neurons in *Cntnap2*-KO mice. It should be noted that none of the ASD mouse models examined by us showed impaired SP behavior, while all of them showed impairments in ESP ^14^.

Overall, our results suggest a pivotal role of fact the mPFC in social discrimination, in general, and in emotion recognition ability, in particular. They also suggest that impaired neural activity of this region, which modifies its synchronization with other social behavior-associated brain regions, such as the PVN and MeAD, is involved in the deficits exhibited by *Cntnap2*-KO mice in terms of emotion recognition. Such impairments in mPFC activity may also underlie similar deficits observed in other murine models of ASD ^14^, suggesting a common brain pathway that integrates the effects of multiple ASD-associated mutations into a specific deficit in emotion recognition.

## Methods

### Animals

#### Source

Adult male and female C57BL/6J mice (C57, 12-14 weeks old) were purchased from Envigo (Rehovot, Israel). *Cntnap2*^−/−^ mice ^17^ (12-14 weeks old) used in this study were maintained by crossing heterozygous mutant males with C57BL/6J females. All *Cntnap2*^−/−^ (KO) and WT mice used in this study were obtained from crossings of heterozygous animals and born with the expected Mendelian frequencies. For generating pure WT mice, WT males and females were crossed to produce pure WT offspring that were later used for behavioral testing at eight weeks of age.

#### Genotyping

Ear tissue samples were collected from offspring mice aged 21 days for genotyping by polymerase chain reaction (PCR) using the following primers:

WT forward primer: TCAGAGTTGATACCCGAGCGCC;
WT reverse primer: TGCTGCTGCCAGCCCAGGAACTGG;
Mutant forward primer: TTGGGTGGAGAGGCTATTCGGCTATG;
Mutant reverse primer: TGCTGCTGCCAGCCCAGGAACTGG.

#### Maintenance

All animals were kept in sex-matched groups of 2-5 mice per cage at the animal facility of the University of Haifa under veterinary supervision, in a 12 h light/12 h dark cycle (lights on at 21:00), with *ad libitum* access to food (standard chow diet, Envigo RMS, Israel) and water.

#### Experiments

Experiments were performed in the dark phase of the dark/light cycle in a sound- and electromagnetic noise-attenuated chamber. All experiments were approved by the Institutional Animal Care and Use Committee of the University of Haifa (Ethical approval #1077U).

#### Ethical statement

All experiments were performed according to the National Institutes of Health guide for the care and use of laboratory animals and approved by the Institutional Animal Care and Use Committee (IACUC) of the University of Haifa.

### Behavioral assays

Male and female KO, WT, and pure WT mice performed multiple social discrimination tasks, as previously described ^14,50,51^. Each session comprised 15 minute window of habituation in the arena with empty triangular chambers in opposing corners of the arena, as previously described ^52^, followed by the subject mice performing the task of interest for 5 minutes (encounter).

In the SP task, subjects were exposed to an adult novel group-housed male mouse (social) and a Lego toy (object), separately located in individual chambers at opposing corners of the arena. In the SxP task, subject mice encountered novel adult male and female stimulus animals. In the SSP task, subject mice encountered a novel stimulus animal stressed by 15 min restraint in a 50 ml Falcon tube with an opening for air, and a non-stressed (neutral) stimulus animal. In the ISP task, subject mice encountered novel group-housed and socially isolated (for 1-2 weeks) stimulus animals. Each isolated stimulus animal was used for two non-consecutive tests. All stimulus animals used in all four tests were novel adult C57BL/6J mice. In the FSI task, subject mice freely interacted with a group-housed, same-sex age-matched novel stimulus animal for 5 min.

### Electrophysiology

#### Electrode array surgery

Mice were anesthetized using isoflurane (induction 3%, 0.5%-0.8% maintenance in 200 mL/min air; SomnoSuite) and placed over a custom-made heating pad (37°C) in a standard stereotaxic device (Kopf Instruments, Tujunga, CA). Two burr holes were drilled to allow placement of ground and reference wires (silver wire, 127 µm, 300-500 Ω; A-M Systems, Carlsborg, WA). Two watch screws (0-80, 1/16”, M1.4) were inserted into the temporal bone to support the electrode array (EAr) with dental cement. Four points (at coordinates: AP = 2 mm, ML= −0.3 mm; AP = 1 mm, ML= −2.3 mm; AP = −2 mm, ML= −2.3 mm; AP = −2 mm, ML= −0.3 mm) were marked over the left hemisphere with a marker. The skull covering these marked coordinates was removed after smoothening the bone with a dental drill, and the exposed brain was kept moist with cold, sterile saline. We custom-designed the Ear ^22^ from 8 to 12 individual 50 µm formvar-insulated tungsten wires (50-150 kΩ, #CFW2032882; California Wire Company) to target the PrL, IL, AcbC, AcbSh, LS, PVN and MeAD. Before implantation, the EAr was dipped in 1,1’-Dioctadecyl-3,3,3’,3’-tetramethylindocarbocyanine perchlorate (42364, Sigma-Aldrich) to visualize electrode locations post-mortem. The reference and ground wires were inserted into their respective burr holes and the EAr was lowered onto the surface of the exposed brain using a motorized manipulator (MP200; Sutter Instruments). The dorsoventral coordinates were estimated using the depth of the electrode targeting the PVN (AP = −1 mm, ML= −0.3 mm), which was lowered slowly to −4.7 mm. The EAr and exposed skull with the screws were secured with dental cement (Enamel plus, Micerium). Mice were sub-cutaneously injected with Baytril (5 mg/kg; Bayer) and Norocarp (5 mg/kg; Carprofen, Norbrook Lab) post-surgery and allowed to recover for three days. After surgery, implanted mice (subjects) were singly housed so as to not disturb the electrode array.

#### Electrophysiological and video recording setup

Following a brief exposure to isoflurane, subjects were attached to a headstage board (RHD 16 ch, #C3334, Intan Technologies) through a custom-made Omnetics to Mill-Max adaptor (Mill-Max model 852-10-100-10-001000). Behavior was recorded above the arena using a monochromatic camera (30 Hz, Flea3 USB3, Flir). Electrophysiological recordings were made with the RHD2000 evaluation system using an ultra-thin SPI interface cable connected to the headstage board. The optic fiber and SPI cable were fed through a motorized commutator (AlphaComm-1, Alpha Omega, Israel) to reduce cable entanglement during the tasks. Electrophysiological recordings (sampled at 20 kHz) were aligned with recorded video using a TTL trigger pulse and by recording camera frame strobes.

#### Electrophysiological recordings

We recorded the behavior and neural activity of 15 KO and 11 C57 adult male mice (Table S1) while targeting distinct brain regions, as described above. We discounted electrodes mistargeted into the dorsomedial hypothalamic nuclei (DMD) and amygdalo-hippocampal region (AhiAL) from further analysis as these regions do not present specific association social discrimination behavior. All stimuli used for the tasks were unfamiliar to the subject mice. Before experiments, the mice were briefly exposed to isoflurane, and the EAr was connected to the evaluation system. Each recording session was divided into two 5 min stages, namely, a baseline period during which time the subject was alone in the arena in the presence of two empty chambers (or no chambers, in the case of the FSI task) and a task (encounter) period when the subject performed the task. Each subject was evaluated over three times (sessions) of each task. The subjects performed the SP and FSI tasks with 10 min intervals separating each of the three rounds. The ISP and SxP tasks were next performed with similar intervals. Four sessions spread six hours apart were recorded daily, two in the morning and two in the afternoon, (see timeline in Fig. 2A). Sessions were excluded from further evaluation only when the headstage detached from the EAr or in the case of a missing video recording from a session. This accounts for the unequal number of sessions and subjects across tasks.

### Optogenetic experiments

#### Surgery

For the optogenetic stimulation experiments, we performed surgeries on C57 and KO male mice (n=6 per group). These mice were anesthetized using isoflurane (induction 3%, 0.5%-0.8% maintenance in 200 mL/min of air; SomnoSuite) and placed under a heating pad attached to stereotaxic apparatus, in a manner similar to the EAr surgeries described above. The mice were then injected with 300 nl of a viral vector encoding an excitatory opsin (ssAAV-1/2-mCaMKIIα-hChR2(H134R)mCherry-WPRE-SV40p(A), produced by Viral Vector core, ETH, Zurich) with a titer of 2.9*10^12^ viral genomes per ml in the left-hemisphere prefrontal cortex (PFC, AP: 1.9 mm, ML: 0.4 mm, DV: 2.2 mm) after craniotomy. An optic fiber (200 um, 0.66 NA, flat bottom, 3 mm long, Doric lenses) was implanted into the PFC (AP: 1.9 mm, ML: 0.4 mm, DV: 2 mm), with the exposed skull along with part of the ferrule holding the optic fiber were stuck together using dental cement (Enamel plus, Micerium). The mice were held in social isolation for recovery three days post-surgery, and then kept for four weeks in group-housing conditions to allow for full expression of the opsin. To mimic the experimental conditions of EAr-implanted mice, mice with optic fiber implants were kept in isolation for three days before the experiments and throughout this period.

#### Behavioral experiments with optogenetic stimulation

Mice undergoing optogenetic stimulation were video-recorded while performing a similar battery of tasks as Ear-implanted mice. Each subject underwent three sessions of the SP, ISP, and SxP tasks, each with a different stimulation protocol, with the order of tests and stimulation protocols being randomized. The stimulation protocols were: (1) no stimulation throughout the encounter period; (2) stimulation at 10 Hz (1 min long, 473 nm, 5 mW, 10 ms pulse) three times, with a 1-min inter-stimulation interval; and (3) similar stimulation as the second protocol, but at 30 Hz. Before the experiments, the mice were lightly anesthetized with isoflurane and connected to the laser (473 nm, model FTEC2471-M75YY0, Blue Sky Research) for light delivery, manually controlled using a pulse stimulator (Master8, AMPI guiding the laser output). The mice were then allowed to habituate to the arena and empty chambers for 15 minutes, after which time actual experiments with stimuli in chambers were recorded for 5 minutes.

### Histology

Subjects were trans-cardially perfused, and their brains were kept cold in 4% paraformaldehyde in PBS for 48 hours. Brains were washed carefully in PBS, sectioned (50 µm) in the horizontal plane with a vibrating blade microtome (VT 1200s, Leica) and collected onto microscope slides. Electrode marks were visualized (DiI coated, Red) against DAPI-stained sections with an epifluorescence microscope (Ti2 eclipse, Nikon). The marks were used to locate the respective brain regions based on the mouse atlas ^53^. In optogenetics stimulation task subjects, mCherry expression and fiber optic location in PFC slides were validated under the epifluorescence microscope using coronal brain slices.

### Data analysis

#### Behavioral analysis

##### Social discrimination (SP, SxP, SSP and ISP tasks)

Subject behavior was tracked using the TrackRodent software, as previously described^51,52,54^.

##### FSI task

###### Markerless pose estimation

DeepLabCut software (v.2.3.5, maDLC) ^23^ was used to track the positions of the subject and stimulus during FSI sessions. The training set included 600 frames from 3 sessions out of a total of 58 sessions (C57 and KO mice). The following body parts were marked for both subject and stimulus in each frame: Ear_left, Ear_right, Nose, Neck, Trunk, Lateral_right, Lateral_left, Tail_base and Tail_mid (Fig. S1A). The model was trained by 2*106 iterations with default parameters (training frames selected by k-means clustering of each of the three videos, trained on 95% of labelled frames, initialized with dlcrnet_ms5, batch size of 8). The model, when tested on the remaining 5% of labeled frames, gave a training error of 3.32 pixels and a test error of 5.43 pixels. Further average root mean square error was less than 4 pixels for all body parts, except for the subject’s nose, which was less visible due to top-view video recordings and electrical cable tether, marked in the labeled frames (Fig. SB).

###### Supervised classification of subject behavior

To identify behaviors across the entire video dataset, we trained supervised random forest classifiers (RF) using a subset of data (5 sessions of 58) from maDLC-analyzed sessions using SimBA’s GUI. Briefly, the sessions were smoothed with a Stravinsky-Golay filter of 100 ms, and outliers were corrected using SimBA’s in-built criterion (1 AU, median of the data). During free interaction sessions, behaviors of interest included *Sniff_StimBody, Sniff_Stim_Anogenital, Being_sniffed, NosetoNose, Aggressive, Moving, Sit_Idle* and *Grooming*. In detail, each of these behaviors was defined as follows:

*Sniff_StimBody:* The subject sniffs the body of the stimulus spanning the whole body, excluding the head and anogenital regions.
*Sniff_Stim_Anogenital:* The subject sniffs the anogenital part of the stimulus body, including any area below the trunk.
*Being_Sniffed:* The stimulus sniffs any part of the subject body, excluding the head.
*NosetoNose:* The subject and the stimulus sniff each other’s facial/head regions.
*Aggressive:* The subject shows aggressive behavior directed towards the stimulus, like biting, tail rattling, trying to mount the stimulus, and holding the stimulus tail with hind paws.
*Moving:* The subject is moving around in the arena, irrespective of the presence of stimulus.
*Sit_Idle:* The subject sits in a non-motile state, irrespective of the stimulus location in the arena.
*Grooming:* The subject grooms or scratches regions of its body and face with either forepaws or hind paws.

An independent RF classifier was trained to classify each of these behaviors. The parameters and the complete RF used to train these classifiers are shared as experimental data. Moreover, classification reports from sklearn.metrics.classification_report were used to quantify and plot precision, recall, and f1 values for each RF classifier. Those behaviors with an RF model that had a f1 value above 0.8 (*Sniff_StimBody, Sniff_Stim_Anogenital, Sit_Idle,* and *Grooming*, Fig. S1G-J) were included for further comparisons between WT and KO subjects. The remaining 53 sessions were analyzed for the RF classifiers of the four behaviors and further analyzed for electrophysiological parameters, similar to the TrackRodent-analyzed data of the SP, EsP, and SxP tasks.

#### Electrophysiological data analysis

##### LFP power

Only brain regions recorded for more than five sessions across at least three mice were analyzed. All signals were analyzed using custom-made codes written in MATLAB 2020a. We excluded signals recorded during 30 seconds around stimulus removal and insertion times to avoid any artifacts due to these actions. The signals were first down-sampled to 5 kHz and low-pass filtered to 300 Hz using a Butterworth filter. The power and time for the different frequencies were estimated using the ‘spectrogram’ function in MATLAB with the following parameters: Discrete prolate spheroidal sequences (dpss) window = 2 s; overlap = 50%; increments = 0.5 Hz; and time bins = 0.5 s. The power of each frequency band (theta: 4-12 Hz and gamma: 30-80 Hz) was averaged for both the baseline and encounter periods (5 min each). Changes in theta (ΔθP) and gamma (ΔγP) power for each brain region were defined as the mean difference in power between the encounter and baseline periods.

##### Coherence

We used the ‘mscohere’ function of MATLAB to estimate coherence values using Welch’s overlapped averaged periodogram method. The magnitude-squared coherence between two signals, x, and y, was defined as follows:

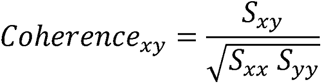

where *S_xy_* is the cross-power spectral density of x and y*, S_xx_* is the power spectral density of x and *S_yy_* is the power spectral density of y. All coherence analysis was quantified between brain region pairs involved in at least five sessions of behavior tasks. Coherence for the baseline period was quantified as the average coherence of all brain region pairs for each task. Changes in coherence (ΔθCo and ΔγCo) during the encounter period between a pair of brain regions were calculated as follows:

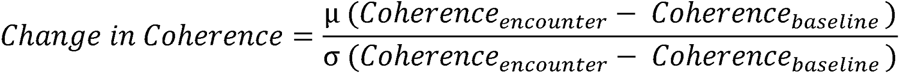

where, Coherence_encounter_ is the absolute coherence value between a pair of regions within a frequency band during a whole encounter period. Coherence_baseline_ is the absolute coherence value between a pair of regions within a frequency band during an entire encounter period.

##### Granger causality

We employed the multi-variate GC toolbox ^55^ to calculate GC values separately for baseline and encounter periods between brain regions for each task and rhythm. For GC analysis, LFP signals were measured at a reduced sampling rate of 500 Hz.

We used the “tsdata_to_infocrit” function to determine the model order of the vector autoregressive (VAR) model. The median model order for all three tasks was 38 (Bayesian information criterion). To further fit the VAR model to our multi-session, multivariate LFP data, the “tsdata_to_var” function of LWR (Levinson-Whittle recursion) in the regression mode, and a median model order of 38 was used separately for the baseline and encounter periods of each task. Next, we estimated the autocovariance sequence of the fitted VAR model with the “var_to_autocov” function. To maximize the computational efficiency of the function, an acmaxlags of 1500 was chosen. This process did not violate the autocovariance VAR model, as was estimated by the “var_info” function. Finally, we calculated the pairwise conditional frequency-domain multivariate GC matrix using the “autocov_to_spwcgc” function and summed the GC for the relevant frequency band (theta or gamma) using the “smvgc_to_mvgc” function.

### Statistical analysis

Statistical analysis was performed using GraphPad Prism 10. Normal distribution of the data was tested using Shapiro-Wilk tests. A paired t-test or Wilcoxon matched-pairs signed rank test was performed to compare various stimuli or conditions for the same group, while a Unpaired t test or Mann-Whitney test was used to compare variables between distinct groups. ANOVA was applied to the dataset when comparing a parameter among multiple groups and *post hoc* multiple comparison tests were performed with Šídák’s correction if a main effect was observed. Similarly, when comparing a parameter that had repeated measurements among multiple groups, repeated measures (RM) ANOVA was performed on the dataset and, in case of a main effect, *post hoc* comparisons between the groups was adjusted using Šídák’s corrections. In the case of GC, multiple Mann-Whitney test results were corrected using FDR correction. When comparing two factors and the interaction between these factors among multiple groups, two-way ANOVA was used, followed with Šídák’s correction of multiple comparisons if a main effect was observed. Similarly, when one of the factors was repeated measurement of the two factors and interaction between them was compared between multiple groups, RM two-way ANOVA was performed. When there was significant main effect, Šídák’s correction of multiple comparisons was performed in *post hoc* tests.

## Statements

### Author contributions

A.N.M.: Formal analysis, Investigation, Methodology, Validation, Visualization, Writing - original draft, and Writing - review & editing; R.J.: Formal analysis, Investigation, Methodology, Validation, Visualization; N.R.; Investigation, Formal analysis; S.N.: Data curation, Project administration, Software, Validation, Visualization, Writing - original draft, and Writing - review & editing. S.W.: Conceptualization, Funding acquisition, Project administration, Resources, Supervision, Writing - original draft, and Writing - review & editing.

## Acknowledgments

We thank Boris Shklyar, Head of the Bio-imaging Unit and Eng. Alex Bizer, the experimental systems engineer of the Faculty of Natural Sciences of the University of Haifa, for his help.

## Supplementary figures

**Figure S1:**
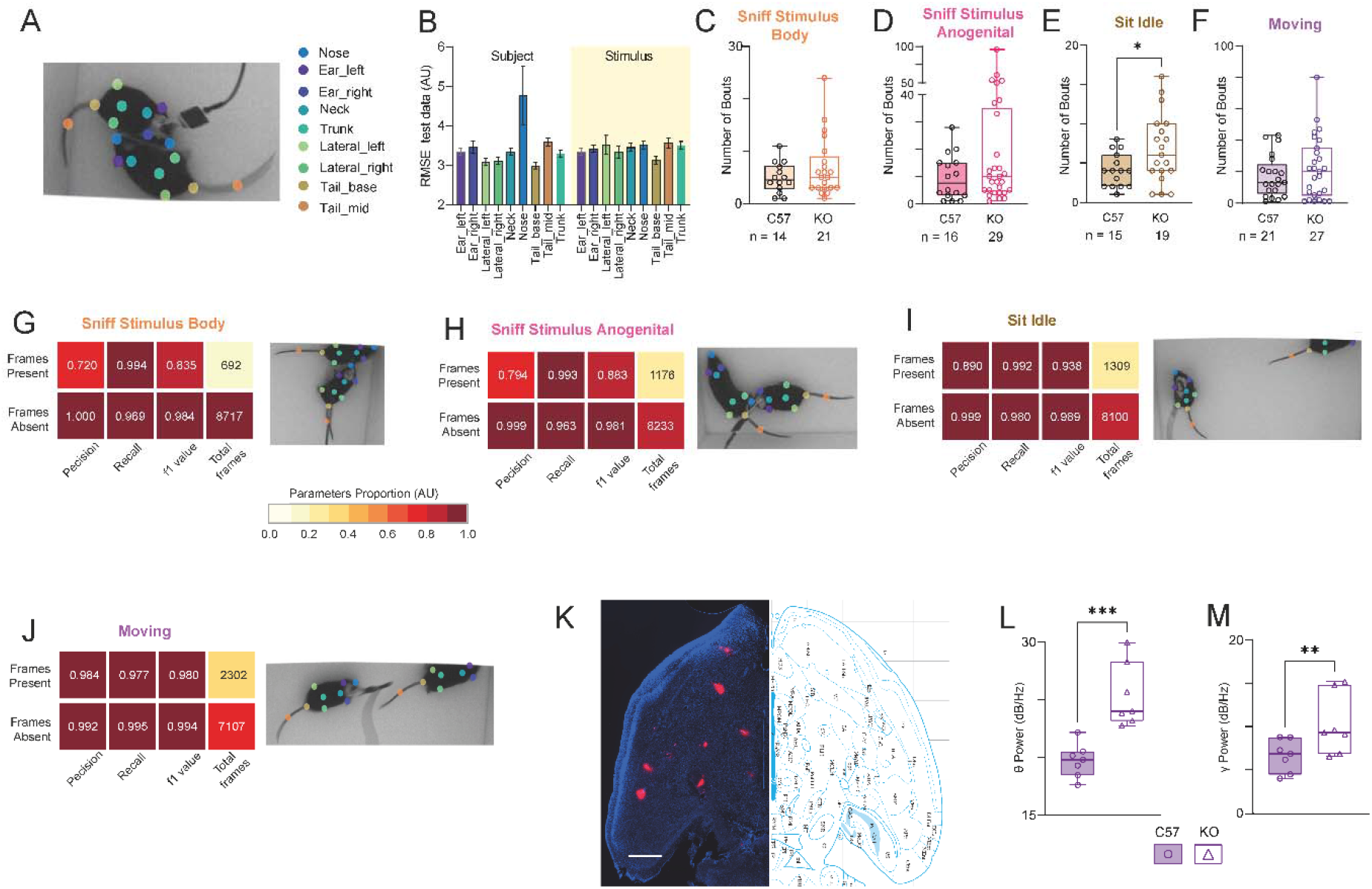
Behavioral annotation using DeepLabCut and SimBA of the FSI task. **A.** Snapshot of a DeepLabCut tracking of a tethered subject interacting with a stimulus animal during a FSI session. Notice the body parts marked by the trained DeepLabCut model and corresponding names of each body part denoted in the right colored labels. **B.** Average (±SEM) root mean square error (RMSE) of the model validated over 5% of all labelled frames. Notice the nose of subject, which is usually obstructed by being tethered, to have poorer error than other body parts. **C.** Median number of *Sniff_StimBody* events (14 C57 and 21 KO sessions annotated with this event; Mann Whitney test, U = 129, P = 0.5529). **D.** As in **C**. for *Sniff_Stim_Anogenital* events (19 C57 and 29 KO; U = 182.5, P = 0.245). **E.** As in **C**. for *Sit_Idle* events (15 C57 and 19 KO; Unpaired t test, t (32) = 2.191, P = 0.0358). **F.** As in **C**. for *Moving* events (21 C57 and 27 KO; U = 257.5, P = 0.5953). **G.** Heat-map of classification metrics from SimBA for presence or absence of *Sniff_StimBody* events in all the frames in which the Random forest model was validated. A snapshot of the SimBA-annotated behavior event is added to the right of the heat-map of the metrics. **H.** As in **G**. for *Sniff_Stim_Anogenital* events. **I.** As in **G**. for *Sit_Idle* events. **J.** As in **G**. for *Moving* events. **K.** Fluorescence image of the horizontal section at −4.7 mm from Bregma of a subject’s left-hemisphere with EAr marks in DiI (red) and gross anatomy depicted by DAPI-stained nuclei (blue). The corresponding panel from the atlas is attached to the right side of the image. Scale = 1 mm. **L.** Median θ power (dB/Hz) of the baseline period from the first session for each subject (Unpaired t test, n = 7 brain regions, t (12) = 4.281, P = 0.0004). **M.** As in **L**. median γ power (t (12) = 2.395, P = 0.0338). Box plot represents 25 to 75 percentiles of the distribution, while the bold line is the median of the distribution. Whiskers represent the smallest and largest values in the distribution. *p<0.05, **p<0.01, ****p<0.0001, Mann Whitney test or Unpaired t test.

**Figure S2:**
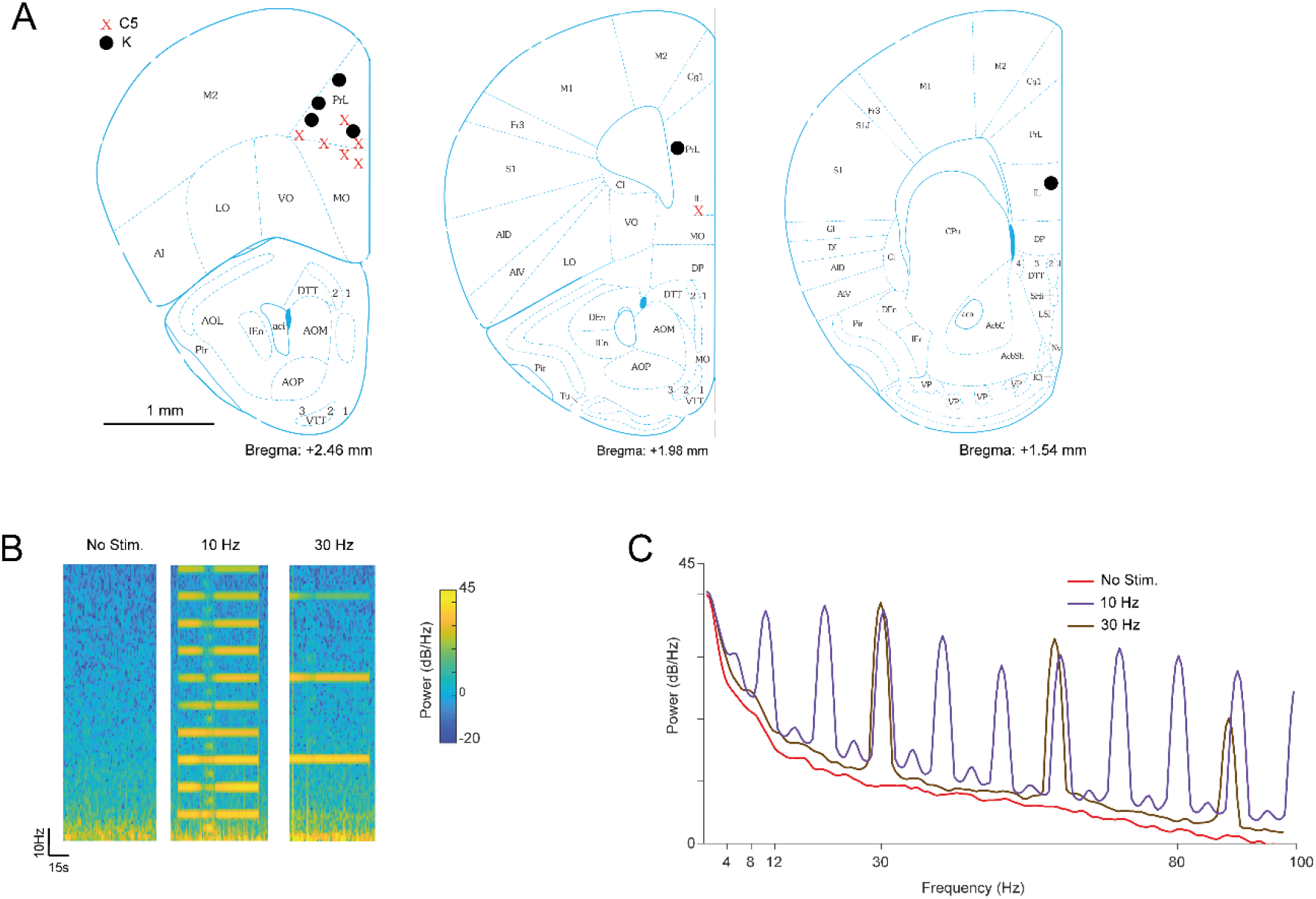
Sites of optic fiber tips and optogenetic stimulation validation. **A.** Coronal brain section panels of the left hemisphere from the mouse atlas at Bregma +2.46, +1.96 and +1.54 with depth of optic fiber marked for C57 (X) and KO (●) subjects. **B.** Spectrograms of recorded mPFC LFP signals for three protocols of optogenetic stimulation in C57 mice expressing ChR2.0 in the left mPFC along with the implanted electrode and optic fiber. The left image corresponds to spectrogram reflecting 1 min of no stimulation from an electrode placed in the mPFC. The middle image is of a 10 Hz (473 nm, 5 mW, 10 ms pulse) stimulation and the right image is of a 30 Hz stimulation. Note the harmonics seen at multiples of the stimulation rate. **C.** Power-spectral density plots of LFP signals from the mPFC electrode shown in B, for no stimulation, and for 10 Hz and 30 Hz stimulation.

